# Mapping the landscape of chromatin dynamics during naïve CD4+ T-cell activation

**DOI:** 10.1101/2020.07.30.228106

**Authors:** Muhammad Munir Iqbal, Michael Serralha, Parwinder Kaur, David Martino

## Abstract

T-cell activation induces context-specific gene expression programs that promote energy generation and biosynthesis, progression through the cell cycle and ultimately cell differentiation. The aim of this study was to apply the omni ATAC-seq method to characterize the landscape of chromatin changes induced by T-cell activation in mature naïve CD4+ T-cells. Using a well-established ex vivo protocol of canonical T-cell receptor signaling, we generated genome-wide chromatin maps of naïve T-cells from pediatric donors in quiescent or recently activated states. We identified thousands of individual chromatin accessibility peaks that are associated with T-cell activation. The majority of these were localized to intronic and intergenic enhancer regions, marked by active histone modifications whilst quiescence was maintained by repressive histone marks. Regions of activation-associated gains in chromatin accessibility were enriched for well-known pioneer transcription factor motifs, and super-enhancer regions associated with distinct gene regulatory networks. These *cis*-regulatory elements together brought about distinct transcriptional signatures in activated cells including TNFa-NFkB signaling, hormone-responsive genes, inflammatory response genes and IL2-STAT5 signaling. Our data provides novel insights into the chromatin dynamics and motif usage of T-cell receptor signaling events in early life. The characterized pathways demonstrate the utility of chromatin profiling techniques applied to bio-banked samples for characterizing gene regulatory elements.

## Introduction

Naïve CD4+ T-cells circulate through the periphery in an actively maintained state of quiescence, ready to mount a robust immune response to pathogens. Quiescent T-cells maintain a tightly condensed chromatin architecture (Rawlings et al. 2010) and cellular program of low energy expenditure whilst surveying for cognate antigen (Wolf et al. 2020), and rapidly undergo substantial re-programming following activation, transitioning toward highly proliferative effector cells. Activation of naïve T-cells initiates rapid functional adaptations which, over the course of days, evolves into heterogenous effector fates with unique helper and regulatory functions with the potential for establishing long-lived memory phenotypes. Activated T-cells rapidly increase nutrient uptake, ramp up translational activity and switch to glycolytic pathways to provide the energy required to support cell growth(Phan et al. 2017), a massive proliferative response and the acquisition of effector functions. These adaptive changes are well understood to be underpinned by epigenetic (Tough et al. 2020), metabolic (Phan et al. 2017), transcriptional and proteomic (Wolf et al. 2020) changes.

At the nuclear level, T-cell receptor (TCR) signaling induces dynamic re-positioning of nucleosomes at promoters and enhancers to allow for transcriptional changes (Schones et al. 2008). These dynamic changes in the chromatin landscape enable interactions between sequence-specific transcription factors (TF) with regulatory DNA elements. Although promoters are the primary sites of transcription initiation, enhancers are major determinants of cell-specific transcriptional and physiological adaptations (Heinz et al. 2015). The assay for transposase-accessible chromatin (ATAC-seq) has gained in popularity as a method to map chromatin accessibility corresponding to TF binding sites and nucleosome positioning (Schep et al. 2015) at the genome-wide scale, due to its high resolution and low cell input, enabling ex vivo analyses (Buenrostro et al. 2013; Scharer et al. 2016). A more recent variant of the ATAC-seq method known as omni-ATAC has demonstrated advantages for removing unwanted mitochondrial reads and exhibits better performance on fixed and flash-frozen material (Corces et al. 2017). ATAC-seq integrated with TF binding motifs has proven increasingly useful for uncovering the dynamic changes in enhancer landscapes and predicting key regulatory events that bring about chromatin remodeling. The dynamic remodeling of enhancer landscapes and differential TF motif usage is a characteristic of distinct T-helper subsets (Bonelli et al. 2014). The majority of data available to date has been performed on neonates (Henriksson et al. 2019), adults (Yukawa et al. 2020; Wolf et al. 2020) or murine cells (Rawlings et al. 2010; Chisolm et al. 2017; Champhekar et al. 2015; Ungerbäck et al. 2018), and there is a paucity of data on infants and young children. Thus, our goal was to examine the utility of omni-ATAC for characterizing chromatin dynamics and inferring gene-regulatory networks in paediatric biobanked samples. We have previously described deficiencies in T-cell activation transcriptional networks and activation-induced regulation of DNA methylation in young infants who developed IgE-mediated food allergy (Martino et al. 2011, 2018). Studying T-cell activation responses at the molecular level has translational potential for understanding disease mechanisms and uncovering novel molecular targets.

In this study we isolated mature naïve T-cells from 6 healthy paediatric donors and studied chromatin dynamics in the canonical T-cell receptor signaling pathway using an identical protocol as published previously by us (Martino et al. 2018). This allowed us to analyze stimulation-dependent chromatin changes in the context of previously collected transcriptomic data. By integrating additional epigenetic data sets we undertook an epigenomic analysis of the paediatric T-cell activation response. Our data are largely consistent with previous studies, demonstrating the utility of omni-ATAC for characterizing the enhancer landscape and motif usage in paediatric bio-banked samples, as a prelude to future studies of disease mechanism.

## Results

### Post-alignment QC

We isolated mature naïve T-cells from 6 healthy infants and studied chromatin dynamics in the canonical T-cell receptor signaling pathway using an identical protocol as published previously by us (Martino et al. 2018). Naïve T-cells were activated with anti-CD3/anti-CD28 beads for 72 hours (activated nCD4T) with matched un-stimulated control condition (quiescent nCD4T). Using cell tracing dyes this protocol results in 3 – 4 distinct T-cell divisions expanding the clonal population on average by 2-fold (expansion index 2.044 [range 1.80 – 2.23], Fig. 1A). After 72 hours all cells were recovered for chromatin profiling. Previous analysis demonstrated activated cells harvested in this phase represent a transitional population of highly proliferative early effector phenotypes (Martino et al. 2018). We generated maps of genome-wide chromatin accessibility to identify epigenomic elements that bring about the stimulation response in resting and activated cells. Post-alignment quality control indicated high mapping efficiency with overall alignment rates 96% or higher. Most reads were enriched at transcriptional start sites for both activated and resting nCD4T. Fragment length distribution plots yielded high resolution of nucleosome-free and nucleosome-occupied reads. Reads were highly enriched at universal DNAse1 hypersensitivity regions identified by the ENCODE consortium (Yue et al. 2014) and enhancer regions indicative of regulatory DNA elements. Activated nCD4T cells exhibited a higher number of reads at promoter regions compared with resting nCD4T (Supplemental Fig. 1).

**Figure 1.**
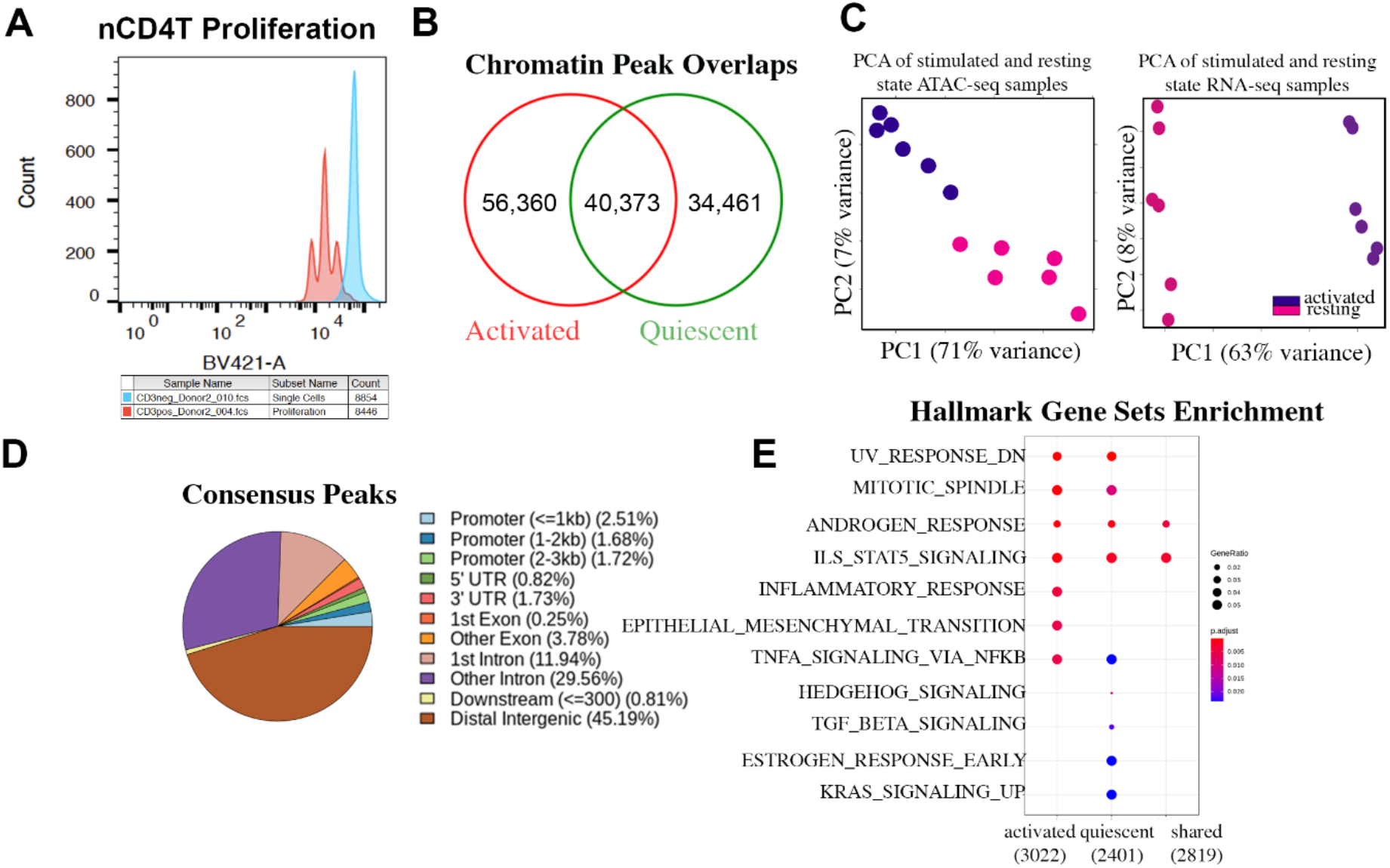
Activation of nCD4T induces widespread changes in chromatin accessibility. **(A)** nCD4T stained with CellTrace Violet and stimulated in culture for 3 days. Discreet peaks represent successive generations of live cells. The unstimulated parent generation is show in blue. **(B)** Venn diagram showing counts of chromatin accessibility peaks. **(C)** Principal component analysis of ATAC-seq peaks (left) and RNA-seq transcripts (right). Each sample is projected into 2D space in a way that best explains variance. **(D)** Annotation of consensus chromatin accessibility peaks to genomic regions of the hg19 genome. **(E)** Gene sets enrichment analysis of molecular signatures enriched in accessibility peaks.

### Open chromatin peak occupancy

Compared to resting nCD4T, the genomes of activated cells were more accessible as evidenced by a larger number of open chromatin peaks. We identified 74,834 consensus peaks in quiescent nCD4T and 96,733 peaks in activated nCD4T, with 40,373 common peaks (Fig. 1B). Principal component analysis of chromatin profiles and publicly available RNA-seq data from our previous study (GSE114064) revealed distinct chromatin signatures for activated and quiescent nCD4T, concomitant with distinct transcriptional programs (Fig. 1C). We annotated consensus peaks to the hg19 reference genome and examined the distribution of peaks across genomic features. The pie chart in Fig. 1D indicated that at least two-thirds of peaks were annotated to enhancer regions (distal intergenic and intronic), with only a small percentage of peaks localized to promoter regions. The latter is consistent with the typical pattern of ATAC peaks representing a mixture of different *cis-*regulatory elements such as enhancers and promotors (Thurman et al. 2012). Hypergeometric testing of peaks revealed quiescent and activated nCD4T shared many regions of open chromatin at genes in the IL2-STAT5 signaling pathway, TNFa-NFkB signal transduction genes and hormone response genes (Fig. 1E). Peaks of chromatin accessibility unique to activated cells were enriched at inflammatory response genes and genes involved in the gain of migratory capacity represented by the ‘epithelial-mesenchymal transition’ pathway. In contrast, peaks of accessibility that were unique to resting nCD4T were enriched in TGF-Beta signaling, estrogen response genes and KRAS signaling (Fig. 1E).

### Differential Binding Analysis

We quantified the number of differentially accessible activation-induced chromatin changes by formally testing MACS2 peaks between quiescent and activated nCD4T. Short-term activation of the T-cell receptor induced substantial changes in the chromatin landscape comprising 43,269 chromatin peaks (q < 5%) that were differentially accessible, of which 5,607 exhibited a minimum absolute log2 fold change in accessibility of +/-2.0 (Fig. 2A). Of the 5,607 differentially accessible peaks, a total of 1,089 peaks gained accessibility in activated nCD4T whilst 4,518 peaks reduced in accessibility (Fig. 2B). Chromatin regions that gained accessibility significantly (FDR < 0.05) coincided with active chromatin marks (H3K4me1, H3K27ac) in primary T-cells, whilst peaks that reduced in accessibility upon activation coincided with repressive marks (H3K27me3 see refs in Wong) (Fig. 2B). We identified putative transcription factor binding sites within regulatory regions that gained accessibility. In total 114 transcription factor motifs were enriched (Fisher’s exact P<0.05) within regions that gained accessibility (Fig. 2C). The most enriched motif was SPIB, a transcriptional activator that acts as a lymphoid specific enhancer to promote the development of IFN-y producing cells (Li et al. 2014). In addition to this novel finding, we identified several well-known factors with a previously described role in TCR signaling and differentiation (STAT5b, JUN, FOS, BACH1, NFATC4, BATF3). This analysis suggested that chromatin landscape changes associated with nCD4T activation prime for differentiation into T-effectors. In support of this, we found that regions of accessible chromatin significantly coincided with DNAseI hotspots found in differentiated T-effectors (Th17, Th1, Th2, Tregs) and were relatively depleted in hotspots unique to naïve precursors and monocytes (as a negative control (Fig. S1)). To corroborate the motif prediction analysis, we used transcription-factor ChlP-seq data available through the ENCODE consortium to test whether regions of that gain accessibility in response to activation coincide with experimentally derived ChIP-seq transcription factor peaks. There were 7 transcription factor ChIP-seq datasets in ENCODE from lymphoblastoid cells lines available for testing (BATF, BHLH, CTCF, JUND, POUF, STAT5A, TCF3), and enrichment analysis indicated all 7 transcription factors signal peaks significantly (Adjusted P < 0.05) coincided with ATAC peaks of accessibility more than expected by chance (Fig. 2D). JUND and STAT5 exhibited the greatest overlap with ATAC accessibility peaks. As expected, when we performed the same enrichment testing on ATAC peaks that lost accessibility, there was no evidence of enrichment (Adjusted P > 0.05, data not shown). We next input the list of 114 identified transcription factors into the GSEA molecular signatures database and performed gene set enrichment analysis, which revealed strong enrichment for the TNFa-NFkB signal transduction pathway (q < 2.68^-06^), UV response (q < 1.92^-02^), KRAS response (q < 2.28^-02^) and Estrogen response (q < 2.28^-02^, Table S1) identified previously as enriched in ATAC peaks (Fig. 1E).

**Figure 2.**
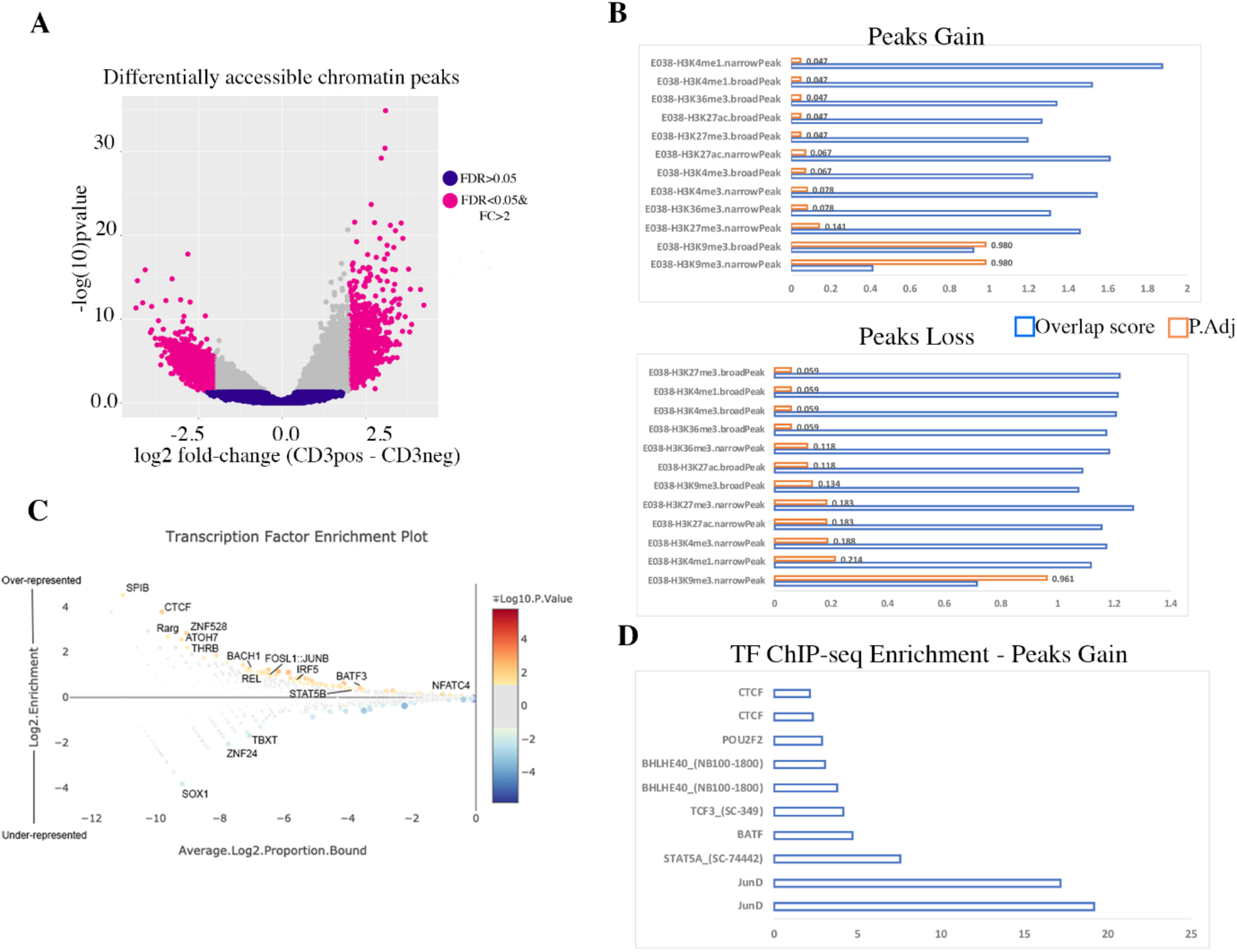
Differential accessibility of chromatin peaks in activated versus resting cells. **(A)** Volcano plot of differentially accessible peaks. Each data point represents a consensus peak. **(B)** Statistical overlap of stimulus-dependent accessible regions with histone ChIP-seq peaks from primary T-cells. **(C)** Enriched transcription factor motifs detected in stimulus-dependent accessible regions. **(D)** Statistical overlap of stimulus-dependent accessible regions with transcription factor ChIP-seq peaks from ENCODE.

### Super-enhancers drive T-lymphoid specific gene regulatory networks

Recent studies suggest core transcription factors can bind clusters of enhancer elements, known as super-enhancers, that can drive interconnected gene regulatory networks (Hnisz et al. 2013). We used the SEanalysis tool to query regions of chromatin accessibility gains against > 330k super-enhancers catalogued from broad H3K27Ac peaks identified from ENCODE CD4+ T-cells. Activation-induced regions that gained chromatin accessibility overlapped 16 known CD4+ T-cell super-enhancers (Table S2). Visualization of ATAC-peaks mapping to the regions identified by the SEanalysis tool showed clear evidence of accessibility gains across the broad region in response to stimulation with concomitant changes in gene expression of nearby transcripts (Fig. 3A and B). These super-enhancer elements formed part of a larger complex gene regulatory network. For example, a super-enhancer annotated to *ARID4B,* a subunit of a co-repressor complex, contains 9 transcription factor binding motifs that were significantly enriched (hypergeometric FDR=2×10^-08^) in interferon alpha/beta signaling, JAK-STAT signaling, interleukin signaling and other pathways (Fig. 3C).

**Figure 3.**
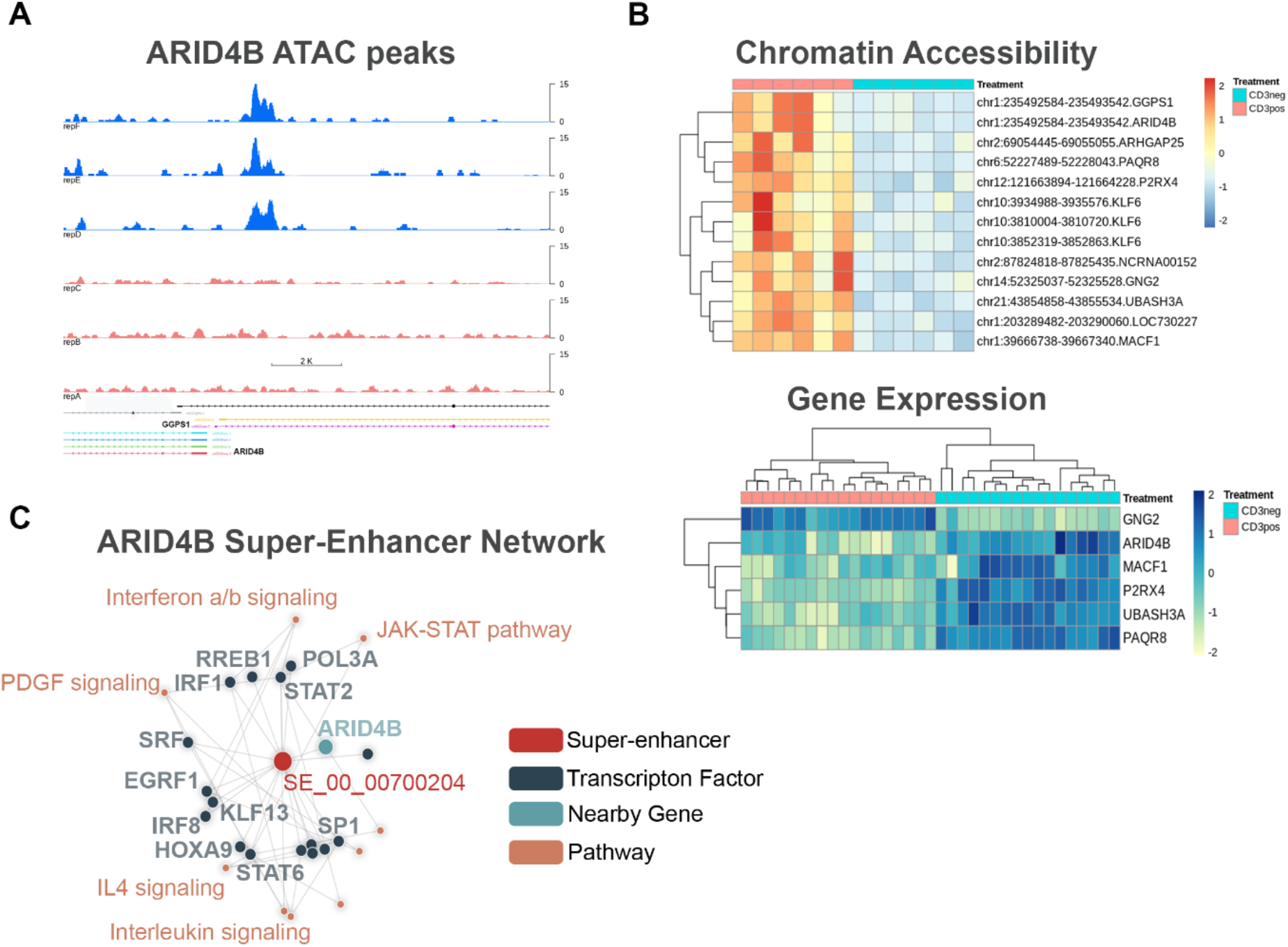
Super enhancer regions detected in stimulus-dependent accessible regions. **(A)** Signal track of ATAC-seq peaks in activated (blue) and quiescent (red) nCD4T. RefSeq transcripts are show in the track below. **(B)** Heatmap of ATAC-seq peaks and RNA-seq transcripts. Rows represent peaks or transcripts and columns represent samples. Cells are colored according to z-score. **(C)** ARID4B superenhancer network diagram showing the regulatory relationship between the identified super-enhancer peak (SE_00_00700204), transcription factor motifs enriched at this peak, and their over-represented down-stream pathways.

### Relationship to transcriptional changes

We next sought to determine how differentially accessible regions associated with activation are related to changes in gene expression. Using transcriptomic data from our previous naïve CD4T study with harmonized laboratory stimulation protocol (GSE114064), we filtered the data set to transcripts overlapping differentially accessible peaks and plotted mean gene expression profiles in activated and quiescent nCD4T. We found that average gene expression levels for transcripts located within differentially accessible regions were broadly similar across treatment conditions (Fig. 4A). Given the high level of enrichment of ATAC-seq peaks in enhancer elements rather than promoters (Fig. 1D), it stood to reason that chromatin changes would likely have more subtle *cis*-regulatory effects on gene expression that would not be obvious as a simple 1: 1 relationship. To explore this further, we first performed a gene sets enrichment analysis on differentially accessible ATAC peaks. Stimulus-responsive regions that gained accessibility were enriched at INFLAMMATORY_RESPONSE and genes in the IL2-STAT5 pathway, whereas those that lost accessibility were enriched in UV RESPONSE, HEDGEHOG and KRAS signaling genes (Fig. 4B). We then tested the hypothesis that expression of these pathways as a whole would differ across treatment conditions. We reasoned that IL2-STAT5 and INFLAMMATORY_RESPONSE signatures would be enriched in activated nCD4T compared with resting nCD4T given the activation-induced gain in accessibility of genes in these pathways. Consistent with this we found strong enrichment (adj. P=0.02 IL2 & adj.P=0.04 INFLAMMATORY) for these molecular signatures in activated nCD4T (Fig. 4C), but there was no evidence of enrichment in resting nCD4T (adj. P = 0.3, IL2 & INFLAMMATORY). Likewise, we found evidence of enrichment transcripts in UV response genes among quiescent nCD4T only but not activated nCD4 T (adj. P=0.03, Fig. 4C).

**Figure 4.**
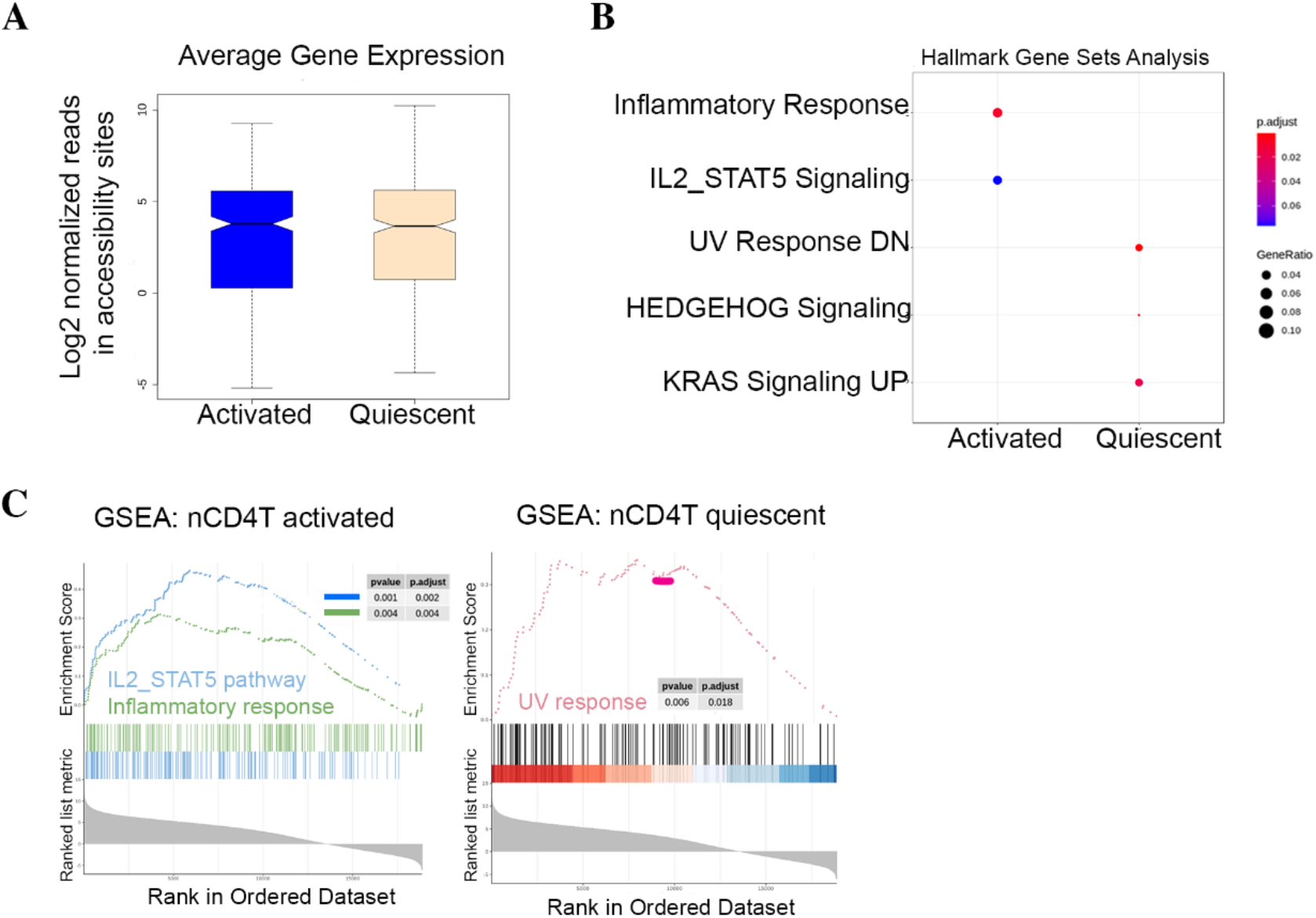
Functional analysis of gene expression at stimulus-dependent accessible regions. **(A)** Boxplot of expressed genes in accessibility sites shows no evidence of global changes in transcriptional output. **(B)** Predicted enriched molecular signatures at differentially accessible peaks (FDR < 0.05 and log2FC +/− 2). **(C)** GenesSets enrichment scores of expressed transcripts validating predicted molecular signatures from ATAC-seq peaks. Top panels show overall enrichment score for the pathway tested. The middle panel tick marks show where the members of the gene set appear in the ranked list of genes. The bottom panel shows the value of the gene’s correlation with phenotype.

Collectively, this analysis suggests that stimulus-dependent chromatin changes drive a broader gene regulatory network comprising both direct and indirect interactions. To illustrate the latter point we identified differentially expressed genes by comparing the transcriptomes of quiescent versus activated nCD4T. In total, 3876 genes were differentially expressed (FDR < 0.05 & logFC > 2). We performed ontology enrichment analysis on the list of differentially expressed genes (DEGs) and differentially accessible regions (DARs). The circos plot in Fig. 5A show the direct overlap between the list of DEGs and DARs was minimal, however functional overlap based on the same gene/region falling into the same ontology term was far more substantial (Fig. 5B). Both differentially accessible regions and genes were enriched among key pathways of T-cell activation including E2F pathway, TNFA_NFKB and IL2_STAT5 signaling, however there were many additional ontologies enriched among differentially expressed genes related to mitotic (MYC targets) and metabolic (MTORC1, Oxidative Phosphorylation) pathways among others.

**Figure 5.**
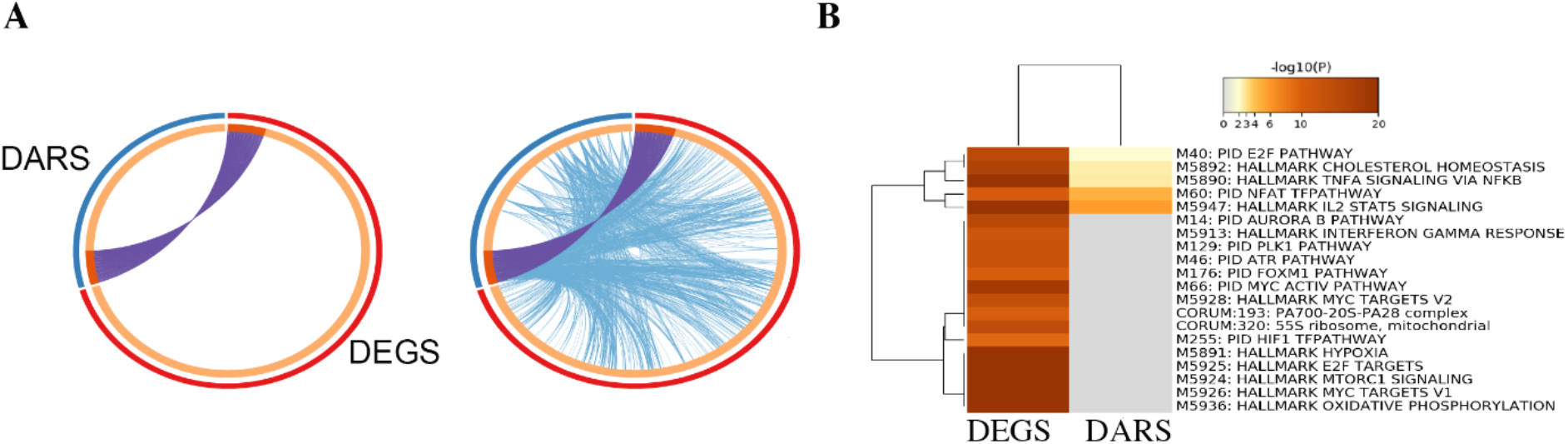
Direct and indirect relationships between stimulus-responsive chromatin peaks and genes. **(A)** Circos plot of differentially accessible regions (DARS: FDR<0.05 & log2FC +/-2) and differentially expressed genes (DEGS: FDR<0.05 & log2C+/-2). Each gene in both lists is represented on the inner arc. Dark orange colour joined by purple chord represents genes that appear in both lists and light orange colour are unique genes. Blue lines link the different genes where they fall into the same statistically enriched ontology term. **(B)** Statistically enriched terms (GO/KEGG terms, canonical pathways, etc.) in chromatin regions and differentially expressed genes. The heatmap cells are colored by their p-values, white cells indicate the lack of enrichment for that term in the corresponding gene list.

## Discussion

T-cell activation induces global remodeling of chromatin accessibility in an orderly and timely manner. These epigenetic changes are coincidental with specific gene regulatory networks that bring about changes in cellular metabolism, proliferative capacity and effector function(Bonelli et al. 2014). In this study we compared genome-wide chromatin accessibility maps between quiescent and activated naïve CD4+ T-cells. Consistent with previous studies (Rawlings et al. 2010) we found that TCR signaling induces wide-spread de-condensation of chromatin, as evidence by substantially higher (~22,000) open chromatin peaks detected in activated cells. The transition to proliferative early effectors appears highly dependent on chromatin remodeling and less so for other epigenetic changes. Our previous genome-wide studies of DNA methylation dynamics in CD4+ T-cell activation revealed no substantial changes in methylation dynamics at 48 hours post-activation (Martino et al. 2012) and 558 CpG dinucleotides that were stimulus-responsive at 72 hours(Martino et al. 2018). This is in contrast to the 43,000 chromatin accessibility changes we detected at the genome-wide level, around 5,000 of which exhibited very large changes. Similarly, Rawlings et al also reported that TCR-induced nuclear de-condensation was not dependent on CpG methylation, nor any substantial net changes in global histone modifications (Rawlings et al. 2010). Chromatin remodeling is therefore a highly dynamic epigenetic mediator of the T-cell activation response.

Chromatin accessibility modulates DNA interactions with transcription factors and the transcriptional machinery. Stimulation associated regions that gained accessibility in our data set were marked with active histone modifications, whilst regions that condense overlapped repressive histone modifications that may serve to suppress alternative cell fates and lineages. In our data set we identified strong enrichment for pioneering transcription factor motifs SPI-1 (also known as PU. 1), CTCF and BATF in regions that gain accessibility. PU.1 is a well-established pioneer factor in early T-lineage commitment that binds gene enhancers (Ungerbäck et al. 2018), supports proliferation and restrains alternative lineages (Champhekar et al. 2015). Binding of SPI-1 can induce chromatin opening and maintain accessibility at target sites. The CTCF DNA binding zinc finger transcription factor plays a spatially organizing role in the genome and promotes precise regulation of developmental gene expression programs. In CD4-T cells, changes in CTCF binding patters are associated with interleukin-2 sensitive metabolic changes (Chisolm et al. 2017). The recruitment of CTCF in T-cells is known to be BATF-dependent, and we detected enrichment for BATF motifs in regions that gain accessibility on activation (Phan et al. 2017).

We also found enrichment of AP-1 family binding motifs (Fos, Jun) which are known pioneer factors that are dramatically up-regulated in response to T-cell activation (Yukawa et al. 2020). Both AP-1 and NFAT (detected in our dataset) are known to play a role in super-enhancer formation in response to T-cell activation (Yukawa et al. 2020). Our analysis identified 13 super-enhancers that were associated with transcriptional changes. Collectively these data characterize the motif usage that bring about the activity change into early effector ‘Th0’ progeny cells. It also highlights the power of the omni-ATAC technique for identifying key regulatory proteins for experimental follow-up with specific transcription-factor ChIP.

The relationship between chromatin landscape dynamics and transcriptional state changes was complex in our interpretation. We found that stimulus-dependent chromatin accessibility changes were enriched in IL2-STAT5 signaling and inflammatory response genes, and these molecular signatures were significantly enriched in activated cells. These are extremely well characterized transcriptional pathways in T-lymphocyte responses (Ross and Cantrell 2018), thus validating the utility of the omni-ATAC technique for deciphering the underlying gene regulatory networks associated with chromatin state changes. It is noteworthy that we did not identify obvious direct relationships between chromatin changes and gene expression, which may be expected given the majority of chromatin remodeling occurred at intronic or distal intergenic sites, suggesting more complex regulation of gene expression. A recent proteomic study has demonstrated that naïve T-cells maintain a reservoir of glycolytic enzymes and un-translated mRNAs that are immediately mobilized in response to activation, allowing naïve cells to kick-start glycolysis and protein synthesis (Wolf et al. 2020). Thus, chromatin responses account for only a portion of the T-cell activation transcriptional response and comprise one of several regulatory mechanisms that underpin T-cell responses. An alternative explanation and potential limitation are that directly relationships were difficult to infer as the gene expression data were from another experiment and therefore may have captured the same biological snapshot albeit at a slightly different time. Although the ex vivo protocols were harmonized, we cannot rule out experimental variation as posing challenges for inferring direct relationships.

The strengths of this study include utilizing a strategy to study canonical TCR signaling by presorting naïve CD4+ T-cells, resulting in a pure and homogenous population. This circumvents difficulties in interpretation posed by co-culture with antigen-presenting cells as chromatin dynamics are likely influenced by secreted factors and cell to cell interactions from accessory cells. By focusing on naïve CD4+ cells rather than total CD4+ cells, which are a mixed population, we avoided confounding due to cellular heterogeneity. We utilized existing RNA-seq data and integrating ChIP-seq and DNAse1-sep datasets from the ENCODE project. These integrations mutually validate the reliability of the ATAC-seq data and aid in biological interpretation (Yan et al. 2020). Caveats include studying only one time-point in the T-cell activation response, and lack of functional data on cytokine responses and cell surface marker changes. This was deemed outside the scope of the current study, as we have characterized these responses in previous studies (Martino et al. 2018)and our focus in the current study was on the chromatin response. We also identified areas of improvement in the omni-ATAC protocol, namely reducing the number of PCR cycles to reduce duplication rates. We were not able to perform a motif foot printing analysis as we did not have sufficient depth of sequencing to accurately call footprints. Overall, this study characterized the chromatin dynamics that bring about the ‘Th0’ early effector progeny and their respective transcriptional state. Importantly, we have done this ex vivo on infant biospecimens demonstrating an approach amenable to paediatric cohort studies. Our future studies will build on the methodology here to study the epigenetic regulation of T-cell activation in disease phenotypes such as allergy and autoimmunity.

## Materials and Methods

### Subject selection

Subjects were recruited through Princess Margaret Hospital in Perth, Western Australia as part of a community-based program of allergy prevention. All subjects used in this study underwent prospective clinical assessments at 1, 2.5 and 5 years of age, including phenotyping for allergic outcomes and general health and donated venous blood for cryopreservation according to institutional ethics committees. Inclusion criteria for selecting biospecimens for this study included equal numbers of males (n=3) and females (n=3), subjects were 1-year of age at time of biospecimen collection, subjects did not receive any interventions, subjects had more than 1 vial of cryopreserved peripheral blood mononuclear cells (PBMC) in the biobank. Exclusion criteria included any congenital malformations, any primary immune deficiency or clinically significant illness that would affect normal hematopoietic development. General characteristics of the cohort are provided in Table S3.

### Isolation, activation and expansion of naïve CD4+ T-cells

Cryopreserved PBMC were thawed in RPMI media (Gibco) supplemented with 10% fetal bovine serum (FBS), Pen-Strep and benzonase (25U/mL) maintained in a 37-degree water bath. After thawing, cells were washed twice, counted and viability checked by trypan blue. Cell recoveries ranged from 8 – 20 million PBMC with viabilities higher than 90%. Naïve CD4+ T-cells (CD3+CD4+CD45RA+CD45RO-) were purified from PBMC using the EasySep Human CD4+ T-cell Isolation Kit (Stemcell Technologies) to >95% purity according to manufacturer’s instructions. Yield of naïve T-cells ranged from 1 – 2.5 million cells. Naïve CD4+T cells were pre-labelled with 5mM CellTrace Violet division tracking dye (Thermo Fisher) according to manufacturer’s instructions and seeded into 96-well polystyrene plates at 80,000 cells per well in RPMI media with 10% FBS, Pen-Strep and human recombinant interleukin-2 (210U/mL, R&D systems). For activation, 2uL of Human T-cell Activator Dynabeads CD3/CD28 (Life Tech) was added to each well reserved for activation, with an equal number of un-activated wells. Cells were incubated for 72 hours at 37 degrees and 5% CO2 before harvesting. At culture end-point, cells were thoroughly resuspended, and beads were removed with replicate wells combined into a single tube for ATAC-seq. A proportion of replicate wells was reserved for proliferation analysis on the BD Fortessa cytometer with 405nm excitation and 450/40 bandpass emission filter.

### Omni ATAC-seq

We employed the omni-ATAC method of Corces. 80,000 viable naïve T-cells were pelleted and lysed in lysis buffer containing 10mM Tris-HCl, 10mM NACl, 3mM MgCL_2_, 0.1% NP40, 0.1% Tween20 and 0.01% Digitonin for 3 minutes on ice. Cells were washed with 1mL of cold wash buffer (lysis buffer without NP40 or Digitonin) and nuclei were pelleted in a centrifuge at 800 RCF for 10 min at 4 degrees. Pelleted nuclei were transposed with Tn5 transposase (Illumina) in TD buffer (Illumina) supplemented with Digitonin (0.1%) and Tween20 (0.01%) for 30min at 37 degrees. Transposed DNA was purified using Zymo DNA Clean and Concentrator-5 Kit (Zymo research) according to manufacturer’s instruction. DNA recoveries were measured on the Qubit fluorometer (Invitrogen). Library amplification was performed using Nextera DNA library prep kit with Nextera Index Kit (Illumina) as per manufacturers instruction. The number of PCR amplification cycles was determined by qRT-PCR using Quanitfast SYBR Green PCR mastermix (Qiagen) and Nextera Primer I5 and I7 Indexes for 5 cycles. The number of additional cycles was determined by a second round of qPCR performed on partially amplified libraries based on the CT value reading taken at 1/3 the fluorescence curve. Two step size selection was performed using AMPure XP beads (Beckman Coulter). Libraries were run on the LabChip GXII fragment analyser and quantitated on the Qubit fluorometer. Libraries were shipped on ice to Novogene (China) for pooling and sequencing on 2 lanes of the Illumina HiSeq at 2×150 paired end reads to generate 50 million reads per sample.

### Bioinformatics

Raw fastq files were analysed using the Multiqc program to generate QC metrics and were processed using the ENCODE official ATAC-seq pipeline version 1.4 specified <u>here</u>. Briefly, adapters detection and trimming were performed using cutadapt (1.91.) and trimmed reads were aligned using the Bowtie2 (2.2.6) aligner. Mapping statistics were generated with SAMtools (1.7) and SAMstats (0.2.1). Post-alignment filtering of duplicates was performed using Picard (1.126) and bedtools (2.26). Aligned reads were shifted +4 bp for the forward strand and −5 bp for the reverse strand. Fragment length statistics were generated using Picard (1.126). Peak calling was conducted using MACSv2 (2.1.0) and blacklisted regions were filtered using bedtools (2.26). Irreproducibility analysis was performed on pseudoreplicates using phantompeakqualtools (1.2.1) and IDR (2.0.4) on 300K MACS2 peaks using a threshold of 0.05. Reads were annotated to ENCODE regions using python scripts and bedtools (2.26).

### Data analysis

All data analyses were conducted in R version 4.0.2. MAC2 peaks were coerced to a peakset object using Diffbind (2.16). Consensus peaksets were derived for activated and quiescent cells defined by presence in more than half the replicates in each group. Peaks were annotated to the hg19 genome using ChIPseeker (1.24). Enrichment analysis of peaks in hallmark genesets was conducted using the clusterProfiler package (3.16). Normalized read counts for consensus peaks were computed for each sample using Diffbind, and differential accessibility between activated and quiescent T-cells was determined using a matched pairs t-test using the edgeR package (3.30). Peaks were declared differentially accessible at the genome-wide level of false discovery rate adjusted P-value < 0.05 and those exhibiting a log 2 fold change of +/-2 or greater were further analysed. Peak signal tracks were generated using the rtracklayer package (1.48). Motif detection analysis was conducted using the CIIIDER tool (Gearing et al. 2019) using the JASPAR core vertebrates 2020 reference database using default parameters. Detection of super enhancers was performed using the SEanalysis tool (Qian et al. 2019) to query accessibility peaks overlapping > 330k super enhancers across 542 cells/tissues annotated in the SEdb database, in Genomic Region Annotation mode using ‘closet active’ gene-SE linking strategy. We restricted the analysis to blood tissue only, and further filtered the results to only primary CD4+ cells. Motif occurrences in constituent enhancers of super enhancers were identified using FIMO (find individual motif occurrences) at p-value threshold of 10^-7^ and enriched pathways were identified using hypergeometric testing at a threshold of FDR-adjusted P-value <0.05. We used the GSuite hyperbrowser program (Simovski et al. 2017) to perform a statistical analysis of over-representation of accessibility peaks with ENCODE datasets. To determine which ENCODE datasets exhibit the strongest similarity to accessibility regions we used the Forbes coefficient to obtain rankings of tracks, and Monte Carlo simulation to provide a statistical assessment of the robustness of the rankings of data tracks, using a null model derived from randomizing the positions of the accessibility regions relative to query tracks. All p-values were adjusted for multiple testing using the Benjamini-Hochberg method. RNAseq data from GSE114064 were downloaded for a subset of age-matched healthy control infants and TMM normalized count data were voom transformed using limma (3.44.3). Differentially expressed genes were declared by comparing transcript expression levels between activated and quiescent T-cells using a matched-pairs t-test (limma) at the genome-wide level of FDR-adjusted P-value < 0.05 and log2 fold change +/− 2 or greater. Overlaps between differentially accessible regions and genes as well as pathways enrichment analysis were computed using the metascape tool under default settings (Zhou et al. 2019). Statistically enriched terms (GO/KEGG terms, canonical pathways, etc.) in chromatin regions and differentially expressed genes were identified and accumulative hypergeometric p-values and enrichment factors were calculated and used for filtering. Remaining significant terms were then hierarchically clustered into a tree based on Kappa-statistical similarities among their gene memberships. Then 0.3 kappa score was applied as the threshold to cast the tree into term clusters. We selected the term with the best p-value within each cluster as its representative term and display them in a heatmap.

## Data Access

The data sets generated for this study are deposited in the Gene Expression Omnibus Repository under accession number GSEXXXX

## Acknowledgements

We wish to acknowledge the contribution of Dr Debbie Palmer, Professor Susan Prescott and the Childhood Allergy and Immunology Research team for their contribution to collection of data and samples pertaining to the cohort studied here.

## Footnotes

Supplemental material is available for this article.

